# Efflux pump antibiotic binding site mutations are associated with azithromycin nonsusceptibility in clinical *Neisseria gonorrhoeae* isolates

**DOI:** 10.1101/2020.06.03.132159

**Authors:** Kevin C Ma, Tatum D Mortimer, Yonatan H Grad

**Affiliations:** Department of Immunology and Infectious Diseases, Harvard T.H. Chan School of Public Health, Boston, USA; Division of Infectious Diseases, Brigham and Women’s Hospital and Harvard Medical School, Boston, USA

## Abstract

Lyu and Moseng et al. used cryo-electron microscopy to characterize key residues involved in drug binding by mosaic-like MtrD efflux pump alleles in *Neisseria gonorrhoeae* (1). Isogenic experiments introducing key MtrD substitutions R714G and K823E increased macrolide MICs, leading the authors to predict that non-mosaic MtrD “gonococcal strains bearing both the *mtrR* promoter and amino acid changes at MtrD positions 714 or 823 could lead to clinically significant levels of Azi nonsusceptibility resistance”. We tested this hypothesis by analyzing a global meta-analysis collection of 4852 *N. gonorrhoeae* genomes (2). In support of their prediction, we identified clinical isolates with novel non-mosaic MtrD drug binding site substitutions across multiple genetic backgrounds associated with elevated azithromycin MICs (Table).

---

Of the 4852 isolates, 12 isolates contained non-synonymous mutations at position R174 to amino acids H, L, or C and 7 isolates contained K823 mutations to E or N in the non-mosaic MtrD background. We did not observe substitutions at binding site residues 174, 669, 821, and 825, in line with the authors’ demonstration that isogenic mutants at these codons had identical or lowered macrolide MICs. The azithromycin geometric mean MICs of the clinical isolates with mutations at R714 and K823 were 1.25 µg/mL and 2.12 µg/mL respectively, both of which are above the CLSI azithromycin non-susceptibility threshold (Figure 1a). There was a significant difference in MIC mean distributions comparing MtrD substitution strains with genetically matched controls (p=0.0008, paired samples Wilcoxon test; Supplementary Table).

**Figure 1.**
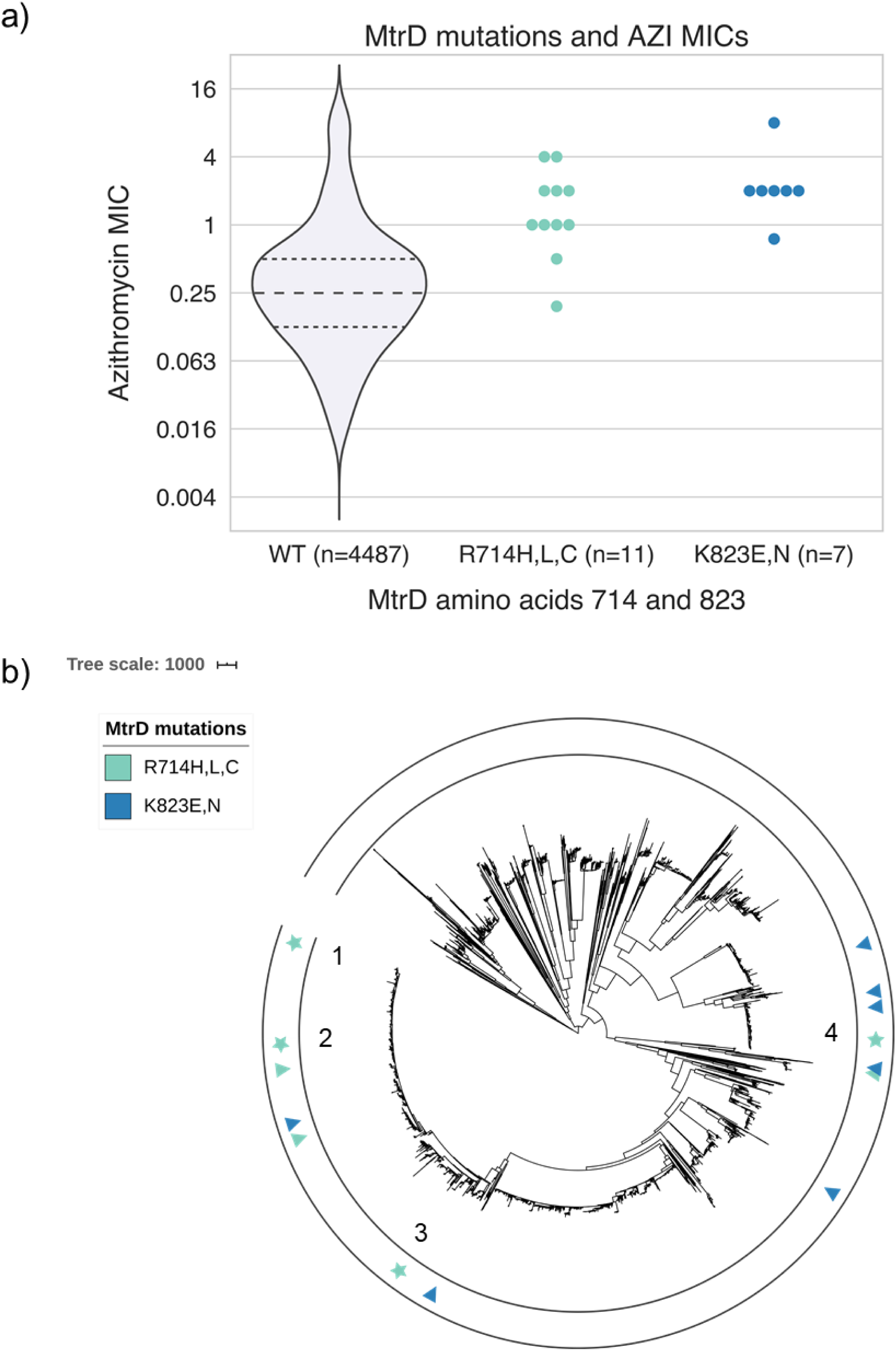
MtrD mutations associated with increased azithromycin MICs have emerged across the *N. gonorrhoeae* phylogeny. a) Comparison of AZI MIC distributions for strains with and without non-mosaic MtrD substitutions at R714 and K823, and b) phylogenetic distribution of MtrD substitution strains in a recombination-corrected phylogeny of the 4852 strains from the global meta-analysis collection. In 1b, triangles indicate singleton strains and stars indicate clusters of two or more strains; cluster number labels correspond to cluster labels in Table.

Nearly all MtrD substitution strains contained *mtrR* promoter mutations that increase MtrCDE pump expression (Table) (3). The isolate with an MtrD R714H mutation and the lowest observed azithromycin MIC of 0.19 µg/mL did not have an *mtrR* promoter mutation, consistent with epistasis across the *mtrRCDE* operon (4). Contributions from ribosomal mutations can also synergistically increase macrolide resistance: the isolate with an MtrD K823E substitution and the highest observed azithromycin MIC of 8.0 µg/mL contained an RplD G70S mutation previously implicated in macrolide resistance (5). Seven MtrD isolates also had mosaic *penA* XXXIV alleles conferring cephalosporin reduced susceptibility, indicating a potential route to dual therapy resistance.

MtrD R714 and MtrD K823 substitutions were each acquired seven times across the phylogeny, suggesting acquisition of the mutation is possible in different genetic backgrounds (Figure 1b). Four of the MtrD K823 acquisitions were associated with more than one isolate descending from the same ancestor, suggesting that these strains are successfully transmitted. In line with this, non-recombinant SNP distances between isolates in each of the four clusters were all below 18 SNPs, with 3/4 clusters below the 10 SNP cutoff previously used as evidence for defining a transmission cluster (6, 7).

Complementing the experimental and structural biology approach taken by Lyu and Moseng et al., we demonstrated using genomics that clinical isolates have acquired novel MtrD binding site mutations which, in combination with *mtrR* promoter and RplD mutations, can result in azithromycin non-susceptibility. As azithromycin resistant strains have been growing in prevalence (8), our data support the inclusion of MtrD binding site residues in future genomic surveillance and genotype-to-phenotype diagnostics and modeling studies for characterizing gonococcal resistance.

## Data availability

All code to replicate analyses is available at https://github.com/gradlab/mtrD-resistance/. An interactive and downloadable version of the phylogeny is hosted at https://itol.embl.de/tree/1281032416307421591107815.

**Table.** MtrD substitution strains, associated metadata, and resistance allele genotypes. Cluster number corresponds to cluster number labels in the Figure 1b phylogeny.

| Accession | Reference | AZI MIC (µg/mL) | MtrD Allele | Cluster | *mtrR* Promoter | RpID G70 Allele | 23S rRNA | PenA Allele |
| --- | --- | --- | --- | --- | --- | --- | --- | --- |
| ERR1469714 | (9) | 1 | R714H | 1 | Adel | WT | WT | XXXIV |
| ERR1528327 | (9) | 1 | R714H | 2 | Adel | WT | WT | XXXIV |
| ERR1514686 | (9) | NA | R714H | 2 | Adel | WT | WT | XXXIV |
| ERR1469709 | (9) | 1 | R714H | 1 | Adel | WT | WT | XXXIV |
| SRR1661243 | (10) | 1 | R714H | 3 | Adel | WT | WT | Non-mosaic |
| SRR2736280 | (11) | 2 | R714H | NA | Adel | WT | WT | XXXIV |
| SRR2736175 | (11) | 2 | R714H | 3 | Adel | WT | WT | Non-mosaic |
| SRR2736167 | (11) | 2 | R714H | 3 | Adel | WT | WT | Non-mosaic |
| ERR349976 | (12) | 0.19 | R714H | NA | WT | WT | WT | Non-mosaic |
| ERR854880 | (13) | 4 | R714L | 4 | Adel | WT | WT | Non-mosaic |
| ERR855125 | (13) | 4 | R714L | 4 | Adel | WT | WT | Non-mosaic |
| ERR855232 | (13) | 0.5 | R714C | NA | Adel | WT | WT | XXXIV |
| ERR363653 | (12) | 0.75 | K823E | NA | Adel | WT | WT | Non-mosaic |
| ERR855395 | (13) | 8 | K823E | NA | Adel | R | WT | XXXIV |
| ERR855128 | (13) | 2 | K823E | NA | Adel | WT | WT | Non-mosaic |
| ERR1067793 | (13) | 2 | K823E | NA | Adel | WT | WT | Non-mosaic |
| SRR2736213 | (11) | 2 | K823E | NA | Adel | S | WT | Non-mosaic |
| SRR2736281 | (11) | 2 | K823E | NA | Adel | WT | WT | Non-mosaic |
| SRR2736124 | (11) | 2 | K823N | NA | Adel | WT | WT | Non-mosaic |

**Supplementary Table 1.** Comparison of AZI MICs of MtrD substitution strains and their nearest neighbors. After log transforming AZI MICs, statistical significance was assessed using a paired samples Wilcoxon test.

**Table – MtrD substitution strains, associated metadata, and resistance allele genotypes.** Cluster number corresponds to cluster number labels in the Figure 1b phylogeny.
| **MtrD substitution strain** | **MtrD Mutation** | **Substitution Strain AZI MIC (µg/mL)** | **Matched MtrD WT strain** | **WT Strain AZI MIC (µg/mL)** | **Non-recombinant SNP distance** |
| --- | --- | --- | --- | --- | --- |
| SRR1661243 | mtrD_714 | 1 | SRR1661226 | 0.25 | 4 |
| SRR2736124 | mtrD_823 | 2 | SRR2736094 | 2 | 7 |
| SRR2736167 | mtrD_714 | 2 | SRR1661226 | 0.25 | 15 |
| SRR2736175 | mtrD_714 | 2 | SRR1661226 | 0.25 | 5 |
| SRR2736213 | mtrD_823 | 2 | DRR124869 | 0.5 | 145 |
| SRR2736280 | mtrD_714 | 2 | ERR855279 | 1 | 17 |
| SRR2736281 | mtrD_823 | 2 | ERR855005 | 0.125 | 12 |
| ERR855128 | mtrD_823 | 2 | ERR1560878 | 0.125 | 106 |
| ERR1067793 | mtrD_823 | 2 | ERR1560869 | 1.5 | 22 |
| ERR855395 | mtrD_823 | 8 | ERR191732 | 1 | 29 |
| ERR855232 | mtrD_714 | 0.5 | ERR854884 | 0.5 | 9 |
| ERR854880 | mtrD_714 | 4 | ERR1471127 | 0.25 | 24 |
| ERR855125 | mtrD_714 | 4 | ERR1560869 | 1.5 | 28 |
| ERR1469709 | mtrD_714 | 1 | ERR1528280 | 0.25 | 11 |
| ERR1469714 | mtrD_714 | 1 | ERR1528280 | 0.25 | 11 |
| ERR1514686 | mtrD_714 | NA | ERR1560893 | 0.5 | 31 |
| ERR1528327 | mtrD_714 | 1 | ERR1560893 | 0.5 | 22 |
| ERR349976 | mtrD_714 | 0.19 | ERR1560863 | 0.38 | 772 |
| ERR363653 | mtrD_823 | 0.75 | ERR363634 | 0.25 | 71 |

## Supplementary methods

### MtrD substitutions

We ran BLASTn on the assemblies using an *mtrD* reference sequence from gonococcal strain FA1090 (Genbank accession: NC_002946.2) and filtered out hits with lower than 99% identity to remove mosaic *mtrD* sequences. LOF alleles were also removed by identifying strains with predicted peptides 90% or shorter than the FA1090 allele. *mtrD* sequences were aligned using MAFFT (version 7.450) (14), translated, and diversity at residues 174, 669, 714, 821, 823, and 825 was characterized using Python (version 3.6.5) and Biopython (version 1.69) (15).

### Nearest neighbor analysis

We calculated non-recombinant SNP distances using snp-dists (version 0.6.3, https://github.com/tseemann/snp-dists) from a recombination masked alignment of polymorphisms produced by a previous analysis of recombination using Gubbins (version 2.3.4) (16). For each strain with a MtrD mutation at codon 714 or 823, we identified the most closely related strain in the dataset without mutations at these positions and with available azithromycin MICs. We used a paired samples Wilcoxon test to test for significant differences between log transformed azithromycin MICs and their nearest neighbor in R (version 3.6.1). Non-recombinant SNP distances were also used to identify genetic distance between MtrD substitution strains within clusters.

